# Lipidomic profiling of plasma extracellular vesicles as an effective means to evaluate the risk of preterm birth

**DOI:** 10.1101/2020.06.07.135764

**Authors:** Qianqian Zhao, Zhen Ma, Xinran Wang, Minling Liang, Wenjing Wang, Fengxia Su, Huanming Yang, Ya Gao, Yan Ren

## Abstract

Preterm birth is the main cause of infant death worldwide and results in a high societal economic burden associated with newborn care. Recent studies have shown that extracellular vesicles play an important role in fetal development during pregnancy. Here, we fully investigated differences in lipids in plasma, microvesicles and exosomes between 27 preterm and 66 full-term pregnant women in the early second trimester (12-24 weeks) using an untargeted lipidomics approach. Independent of other characteristics of samples, we detected 97, 58 and 10 differential features (retention time (RT) and m/z) with identification by multivariate and univariate statistical analyses in plasma, microvesicles and exosomes, respectively. These altered lipids were involved in the formation of the bacterial cell wall and chronic low-level inflammation and oxidative stress. Furthermore, lipids in microvesicles could distinguish patients who experienced preterm labor from controls better than lipids in plasma and exosomes. The candidate lipid biomarkers in microvesicles were also validated by the pseudotargeted lipidomics method. The validation set included 41 preterm and 42 healthy pregnant women. PS (34:0) in microvesicles was able to distinguish preterm birth from healthy pregnancy with higher accuracy. Our study shows that differences in lipids in plasma, microvesicles and exosomes are useful for understanding the underlying mechanisms, early clinical diagnosis and intervention of preterm birth.

## Introduction

Preterm birth, characterized by birth occurring at fewer than 37 weeks gestational age, is a major obstetric health problem leading to poor pregnancy and neonatal outcomes, including neurodevelopmental impairments, hearing and sight problems [1]. Despite remarkable advances in understanding risk factors and mechanisms related to preterm labor and general medical care, the preterm birth rate has not improved. Worldwide, about 15 million infants are preterm each year, and this number is increasing [2]. In the USA, the preterm birth rate has even risen to 12-13% in recent decades, and in Europe and many developed countries, it is generally 5-9% [3]. If women who are at risk of preterm labor are identified early, there would be ample opportunity to adopt appropriate intervention strategies to improve maternal and neonatal outcomes. However, current methods to estimate the risk of preterm birth have low positive predictive value (21% for cervical length and 17% for cervicovaginal fetal fibronectin) and specificity (52% for cervical length) [4]. Therefore, a better understanding of the physiological and pathological mechanisms involved in preterm labor is paramount to more reliably predict and intervene against preterm birth.

Extracellular vesicles (EVs), such as microvesicles and exosomes, were found to play key roles in embryo implantation, pregnancy, and parturition in recent studies [5, 6]. EVs are membrane-enclosed lipid bilayer nanovesicles found in most biological fluids. Microvesicles (MVs) are vesicles 100 nm-1 μm in size that are released from the budding of the plasma membrane, while exosomes (Exos) are 40-120 nm in size and are secreted by exocytotic fusion of multivesicular body with the plasma membrane of cell. EVs contain a diverse array of signaling molecules, including proteins, nucleic acids and lipids [7]. Previous publications have found that the numbers and contents of EVs are significantly different between pregnant and nonpregnant women [8], women at different stages of gestation [9], and women with pregnancy complications and healthy pregnancies [10]. Interestingly, EVs produced by microbes in the genital tract are associated with inflammation and preterm birth [11]. Due to the protective effect of lipid membranes, EV contents are stable in biofluids [12]. Moreover, the levels of contents in EVs are higher than those in originating cells [13]. Thus, EV contents (such as proteins, nucleic acids and lipids) provide a powerful noninvasive method for predicting the risk of preterm birth. Proteins and nucleic acids in EVs related to preterm birth have been characterized [5, 14]. However, the lipids within EVs remain undefined.

Lipids are important structural and functional molecules and are associated with disorders, including metabolic syndrome, inflammation and neurological disorders. Lipids are very diverse in their structures and physicochemical properties, which results in wide variations in biological functions [15]. Lipidomics is a powerful analytical tool to identify and quantify alterations in the lipidome of cells, tissues or body fluids, revealing subtle perturbations caused by diseases, the environment or drugs [16]. The methods of lipidomics analysis include LC-MS, GC-MS and NMR. In practice, LC-MS is frequently used for lipid analysis due to the need to measure lipids with high throughput and reproducibility. The approaches based on LC-MS consist of nontargeted and targeted lipidomic [17]. Recently, a pseudotargeted lipidomics method was developed by Guowang Xu and colleagues [18]. Therefore, the study of lipidomic changes in EVs or plasma related to preterm birth would allow us to discover potential biomarkers for earlier medical diagnoses using LC-MS.

Here, we first used an untargeted lipidomics approach to comprehensively investigate the differences in lipids between preterm birth and healthy pregnancy from plasma, MVs and exosomes in the early second trimester (12-24 weeks). We show that our method is robust and reliable, making it a suitable approach for EV and plasma lipid biomarker discovery. We found that lipids in MVs could distinguish patients who experienced preterm labor from the controls better than lipids in plasma and exosomes. Furthermore, we validated candidate lipid biomarkers in MVs by pseudotargeted lipidomics.

## Materials and Methods

### Materials

LC-MS-grade isopropanol (IPA), methanol (MeOH), and acetonitrile (ACN) were purchased from Fisher Scientific, Inc. (Rockford, IL). Ammonium formate (NH4HCO2) was purchased from Sigma Aldrich (St. Louis, MO), and formic acid (HCOOH) was obtained from Dikma Technologies, Inc. (Beijing, China). Ultrapure water was produced from a Milli-Q system (Millipore, Billerica, MA). Anti-TSG 101 antibody, anti-CD9 antibody, and anti-ALIX antibody were purchased from Abcam plc (Shanghai, China). Goat anti-rabbit IgG (H+L)-HRP was purchased from Beijing Protein Innovation (Beijing, China). SPLASH LipidoMix™ Internal Standard including 15:0-18:1(d7) PC, 15:0-18:1(d7) PE, 15:0-18:1(d7) PS (Na Salt), 15:0-18:1(d7) PG (Na Salt), 15:0-18:1(d7) PI (NH4 Salt), 15:0-18:1(d7) PA (Na Salt), 18:1(d7) Lyso PC, 18:1(d7) Lyso PE, 18:1(d7) Chol Ester, 18:1(d7) MAG, 15:0-18:1(d7) DAG, 15:0-18:1(d7)-15:0 TAG, d18:1-18:1(d9) SM, and Cholesterol (d7) was purchased from Avanti Polar Lipids (Alabaster, AL).

### Specimens

Samples and clinical information were reviewed and approved by the Institutional Review Board on Bioethics and Biosafety, BGI Shenzhen. After obtaining informed consent, a total of 176 maternal K_2_-EDTA plasma samples (12-24 weeks gestation) were collected from the samples remaining after noninvasive prenatal testing. The discovery set consisted of 27 preterm pregnant women (between 32^0/7^ weeks and 36^6/7^ weeks) and 66 full-term pregnant women (≥ 37^0/7^ weeks), who were recruited between February 2013 and March 2016. The validation set consisted of 41 preterm pregnant women (between 32^0/7^ weeks and 36^6/7^ weeks) and 42 full-term pregnant women (≥ 37^0/7^ weeks), who were recruited between January 2018 and May 2018. Gestational age was confirmed based on the last menstrual period date and ultrasound scanning. Maternal age, fetal sex, and birth weight were collected. All subjects had spontaneous labor and had no other obstetric diseases. Detailed information is shown in Table 1. All plasma samples were stored at −80°C.

**Table 1.** Detailed information of preterm birth vs term control pregnancies.

| Characteristic | Discovery Set |  | Validation Set |  |
| --- | --- | --- | --- | --- |
|  | Preterm Birth (n=27) | Control (n=66) | Preterm Birth (n=41) | Control (n=42) |
| Maternal age (Y) (Mean±SD) | 32±5 | 34±6 | 30±5 | 31±4 |
| Gestational age at sample collection(W) (Mean±SD) | 17.9±2.1 | 19.2±2.9 | 17.4±2.5 | 17.6±2.7 |
| Gestational age at delivery (W) (Mean±SD) | 35.2±1.5 | 39.7±0.7 | 35.3±1.2 | 39.6±0.9 |
| Fetal sex(n) (F/M/T) | 13/12/2 | 29/32/5 | 21/17/3 | 24/17/1 |
| Birth weight (g) (Mean±SD) | 2601.9±672.7 | 3594.6±543.7 | 2662.6±812.3 | 3291.0±464.7 |

Y, Year; SD, Standard Deviation; W, week; F, Female; M, Male; T, Twins;

### Extracellular Vesicle Isolation

Extracellular vesicles were isolated by serial centrifugation as previously described with the following changes [19]. Briefly, 250 μl plasma was first diluted to a volume of 750 μl using phosphate-buffered saline (PBS). The diluted plasma was centrifuged (2000 g, 4°C, 10 min) to remove the cell debris. Then the supernatant was centrifuged (8000 g, 4°C, 20 min), discarding the pellets. Next, the supernatant was centrifuged (20000 g, 4°C, 1 h). Pellets were washed with 4°C cold PBS and centrifuged again (20000 g, 4°C, 1 h); the pellets were composed of microvesicles and were collected in 100 μl PBS. The exosomes present in the supernatant were filtered through a 0.22-μm membrane (Millipore) to a final volume of 1 ml and then further centrifuged (120000 g, 4°C, 2 h) in a TLA-120.2 rotor (Beckman Coulter, Brea, CA). Pellets were washed twice with 4°C cold PBS and centrifuged (120000 g, 4°C, 2 h). Finally, pellets were composed of exosomes and were resuspended in 200 μl PBS. All purified microvesicles and exosomes were stored at −80°C.

### Electron Microscopy

The purified exosomes suspended in PBS were dropped onto formvar/carbon film coated copper grids and dried at room temperature. The grids were then washed twice with ultrapure water and stained with 1% uranyl acetate. The grids were air-dried overnight and finally imaged by a JEM-1230 transmission electron microscope (Jeol Ltd., Tokyo, Japan).

### Nanoparticle Tracking Analysis

The size and number of extracellular vesicles (including microvesicles and exosomes) were determined using a NanoSight NS300 with particle tracking analysis (Malvern Instrument, UK). Extracellular vesicles were diluted with PBS, and the measurement was performed according to the manufacturer’s instructions. Each sample was measured three times. The data were analyzed using NTA 3.2 software.

### Protein Quantification and Western Blot

The isolated extracellular vesicles were lysed by repeated freeze-thaw cycles as described previously [20]. Extracellular vesicles were frozen in liquid nitrogen (30 s) five times and −20°C (1 h) two times. Each thawing was performed by sonication for 10 min in a water bath. Protein quantification was determined using the Bradford method [21]. The protein concentration of each sample was measured in duplicate. Proteins were separated by 10% SDS-PAGE and then transferred to a PVDF membrane [22]. For Western blotting, membranes were blocked with 5% nonfat dry milk in TBST (TBS with 0.1% Tween 20) at room temperature for 1 h and hybridized with the primary antibodies at dilutions recommended by the suppliers at 4°C overnight. After washing, the membranes were further hybridized with HRP-conjugated secondary antibodies at room temperature for 1 h. After washing with TBST buffer, the blots on the PVDF membranes were developed with ECL detection reagents. TSG101, Alix, and CD9 were used as primary antibodies [23].

### Lipid Extraction

The lipid extraction method for plasma and extracellular vesicles followed a previous report with minor modifications [24]. Forty microliters of plasma, 100 μl of microvesicles and 200 μl of exosomes suspended in PBS were precipitated by adding 3 volumes of pre-chilled IPA. To provide accurate quantitation of each lipid species, 1.2 μl and 4 μl of a lipid internal standard mixture were added to plasma and extracellular vesicles, respectively. Samples were vortex mixed for 1 min and then incubated at room temperature for 10 min. Subsequently, samples were stored at −20° C to improve protein precipitation and then centrifuged at 14000 g for 20 min. For plasma lipid extracts, the supernatant was further diluted with IPA/ACN/H2O (2/1/1 v/v/v). For extracellular vesicle lipid extracts, the supernatant was dried, and the pellets were dissolved in IPA/ACN/H2O (2/1/1 v/v/v). The samples were stored at −80° C until LC-MS analysis. Equal volumes of lipid extracts of plasma, microvesicles and exosomes were pooled as quality control (QC) samples for monitoring the performance of LC-MS system, respectively [25].

### LC-MS/MS Analysis

For nontargeted lipidomics profiling, an ACQUITY UPLC system (Waters, Manchester, UK) coupled with an electrospray ionization (ESI) source with a G2-XS QTOF mass spectrometer (Waters) was used. The LC-MS/MS methods were used, as described in earlier reports [26]. The LC conditions were as follows: ACQUITY UPLC CSH C18 column (2.1 × 100 mm, 1.7 μm, Waters); mobile phase, (A) ACN/H2O (60/40, v/v) (containing 10 mM NH4HCO2 and 0.1% HCOOH) and (B) IPA/ACN (90/10, v/v) (containing 10 mM NH4HCO2 and 0.1% HCOOH). The LC gradient used was as follows: 0-2 min, 40-43% B, Curve, 6; 2-2.1 min, 43-50% B, Curve, 1; 2.1-7 min, 50-54% B, Curve, 6; 7-7.1 min, 54-70% B, Curve, 1; 7.1-13 min, 70-99% B, Curve, 6; 13-13.1 min, 99-40% B, Curve, 1; and 13.1-15 min, 40-40% B, Curve, 6. The volume of injection was 5 μl. The flow rate was 0.4 ml/min, and the column temperature was maintained at 55°C. The mass data were collected in both positive and negative Centroid MS^E^ mode with an acquisition time of 0.2 s per scan. The low collision energy was set at 6 V, while the high collision energy was set from 19 V to 45 V. The capillary voltage was set at 3 V and 2 V in positive and negative modes, respectively. The cone voltage was set at 30 V in both modes. The source temperature was set at 120°C. The desolvation temperature and gas flow were 450°C and 800 L/h, respectively. Acquisition was performed from m/z 100 to 2000. Leucine enkephalin (m/z 556.2771 in ESI+, m/z 554.2615 in ESI-) was used for lock mass correction, and sodium formate solution was used for mass calibration. Moreover, QC samples were inserted into the analysis sequence at regular intervals to ensure the LC-MS system stability during acquisition.

In the validation experiment, an ACQUITY UPLC system (Waters) coupled with a Q-Trap 6500 MS system (AB SCIEX, Redwood City, CA) was used for HPLC/QQQ MRM MS-based pseudotargeted lipid analysis [18]. The LC conditions were the same as those in the nontargeted lipidomics profiling. The mass data were acquired in positive mode. The MS parameters were as follows: curtain gas was set at 40, and collision gas was set at medium. IonSource Voltage was set at 5500 V, the temperature was set at 500°C, and both Ion Source Gas 1 (GS1) and Ion Source Gas 2 (GS2) were set at 40. The declustering potential (DP) and collision energy (CE) of targeted lipids were determined based on the lipid internal standard mixture. QC samples and lipid internal standards were used to monitor the overall quality of the sample extraction and LC-MS analysis.

### Data processing

For untargeted lipidomics analysis, the MS data files were preprocessed using Progenesis QI software (including peak alignment and picking). Then, the peak table files were processed using metaX software [27]. Features were excluded if they were detected in < 50% of the QC samples or < 20% of the experimental samples. After data filtering, k-nearest neighbor (KNN) was implemented to perform missing value imputation. To correct signal drift, QC-robust spline batch correction (QC-RSC) [25] was performed. After probabilistic quotient normalization, features with a coefficient of variation (CV) ≥ 30% in the QC samples were removed. The filtered data were further used for the following statistical analysis. Lipid identification was based on both parent mass and theoretical fragment. All lipids were matched in three databases, the Lipid Maps [28], the Human Metabolome Database (HMDB) [29] and the Kyoto Encyclopedia of Genes and Genomes (KEGG) [30] with ppm < 10. All identifications were further confirmed manually. The pseudotargeted lipids were processed using MultiQuant software to produce peak areas.

### Statistical Analysis

Univariate and multivariate statistical analyses were carried out by metaX software [27]. Principal component analysis (PCA) and partial least square discriminant analysis (PLS-DA) were employed for sample overview and classification. The PLS-DA models were validated by permutation tests (200 times); fitted models were considered significant if the R^2^ and Q^2^ of the PLS-DA were positive. The variable importance in projection (VIP) score was used to visualize the influence of a variable in the model [31]. Significant differences between preterm birth groups and the control were tested using the two-tailed unpaired t test, and the false discovery rate (FDR) of the p-values was corrected by using the Benjamini-Hochberg FDR algorithm [32]. Potential biomarkers were determined according to VIP ≥ 1, fold change ? 1.2 or ≤ 0.8, and Q value < 0.05. To prevent results from being skewed by protein concentration, gestational age at sample collection, maternal age, fetal sex and birth weight, correlation analysis and multiple linear regression were used. Logistic regression was used to combine the predictive ability of candidate lipid biomarkers, and the receiver operating characteristic (ROC) curve and the area under the curve (AUC) analysis were implemented by SPSS.

## Results

### Characterization of extracellular vesicles extracted from plasma

To establish a procedure for EV lipidomics profiling, we purified MVs and Exos from 250 μl pooled plasma of pregnant women via differential speed centrifugation. Isolated EVs were characterized by transmission electron microscopy, Western blotting and nanoparticle tracking analysis. The morphology of a single exosome observed under a transmission electron microscope is shown in Fig. 1A and was consistent with that described in other reports [33]. Western blotting was used to evaluate the enrichment of exosomes based on centrifuge isolation by detecting the special protein markers of exosomes. As shown in Fig. 1B, the exosome-enriched proteins of Alix, TSG101 and CD9 were found with higher abundance in Exos under the same loading amount of proteins in plasma, MVs and Exos. In the Nanosight determination of vesicles particle size and concentration, as shown in Fig. 1C, the mode size of MVs (93 nm) was markedly larger than that of Exos (60 nm); however, the concentration of MVs (2.3×10^11^ particles/ml plasma) was much lower than that of Exos (3.3×10^12^ particles/ml plasma). These observations indicated that the quality of purified EVs from plasma was good enough for the following lipidomics profiling study.

**Figure 1.**
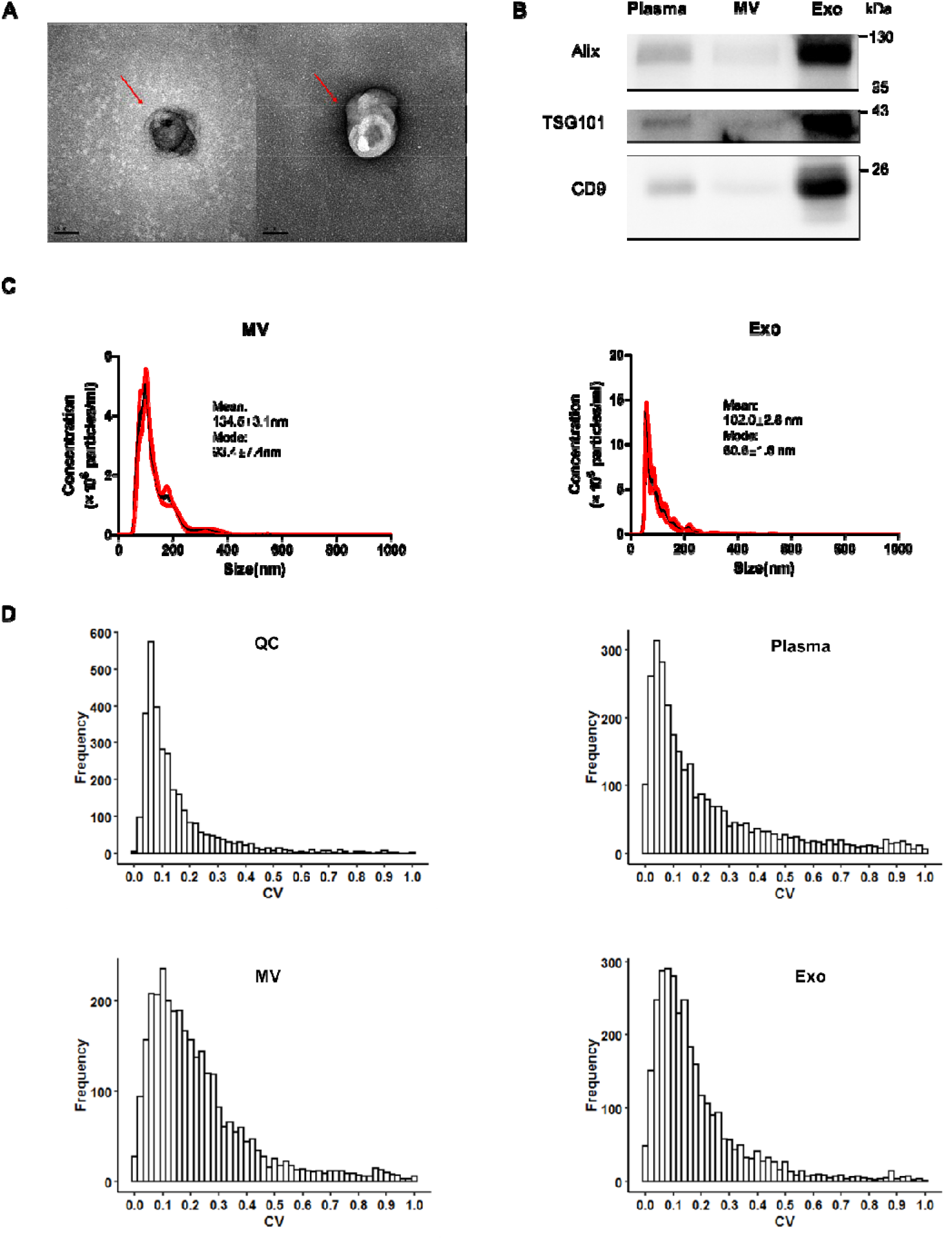
Characterization of extracellular vesicles and repeatability of untargeted lipidomics analytical workflow. A. Representative transmission electron microscope images of isolated exosomes from plasma. Scale bar, 50 nm. B. Western blot analysis of Alix, TSG101 and CD9 in plasma, microvesicles and exosomes. C. Size distribution and particle concentration of microvesicles and exosomes. D. Histograms of the CVs of QC, plasma, microvesicle and exosome samples. MV, microvesicle; Exo, exosome; QC, quality control.

### Reproducibility evaluation of lipidomic profiling from plasma, microvesicle and exosome fractions

As the stability of the lipidomic profiling system is very important for the relative quantification of spectra among different samples, it is necessary to assess the reproducibility of the whole analytical workflow in this study. We performed three technical replications with pooled plasma from pregnant women to evaluate the reproducibility of the lipidomic profiling of plasma, MVs and Exos in positive mode. QC samples (pooled lipid extracts of plasma, MVs and Exos; n=9) were included in the analysis sequence. The CV distribution for all the detected features showed that 88.9% of the features detected in QC samples had a CV < 30% (Fig. 1D), which indicated that the instrument was running in good condition. The percentage of features with a CV < 30% detected in plasma, MVs and Exos was 75.6%, 76.2% and 83.5%, respectively. The results suggest that the LC-MS procedure for our lipidomics profiling analysis is reliable.

### Lipidomic profiling differences in plasma, microvesicles and exosomes between preterm birth and full-term pregnancies

To investigate differential lipids in plasma, MVs and Exos between preterm birth and full-term pregnancies, we collected 27 plasma samples from pregnant women with premature birth and 66 plasma samples from full-term pregnant women. The details of the information are shown in Table 1. It was noted that MVs and Exos were isolated from the same volume of plasma (250 μl). To assess the quality of the lipid extraction and LC-MS analysis, the CVs of lipid internal standards, the total ion chromatography (TIC) of QC samples and PCA with QC samples were assessed. The percentage of lipid internal standards with CV < 20% in both positive and negative modes after pooling into plasma, MVs and Exos was 69.2%, 61.5% and 69.2%, respectively (Table S1). The TIC of QC samples overlapped very well, and the PCA plots showed that QC samples were clustered together (Fig. S1&S2). The results indicated that the data quality met the requirements for the subsequent statistical analysis.

We first applied PCA to reveal discriminant features between the preterm birth and control conditions. The PCA with MV features in positive mode showed better separation between these two groups than the others (Fig. S3A, S3E, S3I). We further applied PLS-DA to search for different features between these two groups (Fig. S3B, S3F, S3J). As shown in Fig. S3C, S3G, and S3K, these models were evaluated by 200 permutation tests. The model goodness of fit (R^2^) and predictive ability (Q^2^) were determined. These models were robust in the plasma and Exos in both positive and negative modes (plasma: positive, R^2^, 0.6449; Q^2^, 0.0369; negative R^2^, 0.5754; Q^2^, 0.0022; Exos: positive, R^2^, 0.596; Q^2^, 0.0368; negative R^2^, 0.5761; Q^2^, 0.109;) and in the MVs in positive mode (R^2^, 0.4785; Q^2^, 0.1672). However, the model was overfitted in the MVs in negative mode (R^2^, 0.4547; Q^2^, −0.2045). The model was not used in the following analysis.

To determine features with significant differences between the preterm birth and control conditions, we adopted the following criteria: VIP ≥ 1, fold change ≥ 1.2 or ≤0.8, and Q value < 0.05. As shown in Table 2, we discovered 330, 632 and 88 features with significant differences between the two groups from plasma, MVs and Exos, respectively. We also employed volcano plots to graphically display the differential features (Fig. S3D, S3H, S3L). We excluded the possibility that other characteristics of patients and protein concentrations of plasma, MVs and Exos (Table S2) skewed the experimental results by using correlation analysis and multiple linear regression. After depletion, 293, 144 and 46 features were retained in plasma, MVs and Exos, respectively (Table 2). The identification of these features was performed by Progenesis QI software against three databases (including Lipid Maps, HMDB and KEGG) and was then further confirmed manually. The features with multiple isomers were identified as the main class of lipids. In total, 97, 58 and 10 features were identified in plasma, MVs and Exos, respectively (Table 2). Subsequently, these differential lipids between the preterm birth and control conditions were classified by the lipid classification system of Lipid Maps (compounds excluded from Lipid Maps were classified by the HMDB classification system). As shown in Fig. 2A, 2C, and 2E, the major types of lipids identified belonged to glycerophospholipids regardless of whether they were identified in plasma, MVs or Exos. However, the detailed compositions of these lipids were different in plasma, MVs and Exos. The differences in the levels of lipid species are shown in Fig. 2B, 2D, and 2E. The trends of these lipid species in plasma are summarized as follows: 1). The following lipids were higher or lower in the preterm birth condition compared to the full-term control condition: PC, PE, PA, PS, PI, PG, and ST; 2). The following lipids were higher in the preterm birth condition than in the full-term control condition: ox-PL, PGP, other-PLs, CDP-DG, AGSL, STC, BAD, SST, FAC, FAld, pPr, IPR, and carbonyl compounds. Interestingly, we also found that undecaprenol (included in pPr) and GDP-D-glycero-alpha-D-manno-heptose levels were higher in the preterm birth condition than in the full-term control condition; and 3). The following lipids were lower in the preterm birth condition than in the full-term control condition: Cer, n-GSL, GSLs, TG, and DG. The trends of these lipid species in MVs are summarized as follows: 1). The following lipids were higher or lower in the preterm birth condition than in the full-term control condition: PG, PS, PE, PI, and n-GSL; 2). The following lipids were higher in the preterm birth condition than in the full-term control condition: PA, ox-PL, MG, DG, SM, Cer, SPH, other SP, SST, STC, FAC, eicosanoids, FAM, pPr, alcohols and polyols. It was noted that undecaprenol (included in pPr) was also higher in the preterm birth condition than in the full-term control condition; and 3). The following lipids were lower in the preterm birth condition than in the full-term control condition: PC and AGSL. The trends of these lipid species in Exos are summarized as follows: 1). The following lipids were higher in the preterm birth condition than in the full-term control condition: PC, PE, ox-PL, and PA. We also detected cotinine glucuronide, which was higher in the preterm birth condition than in the full-term control condition and 2). The following lipids were lower in the preterm birth condition than in the full-term control condition: CL, TG, DG, FE, and Cer. We also investigated the changes in LPs between these two groups and found that all LPs were higher in the preterm birth condition than in the full-term control condition, regardless of whether they were in plasma, MVs or Exos. Detailed information on the differential features with identification is shown in Table S3. Together, these results suggest that differential lipid compositions between preterm birth and control conditions in plasma, MVs and Exos show subtle differences, which is helpful for fully understanding the role of lipids in the pathogenesis of preterm labor.

**Figure 2.**
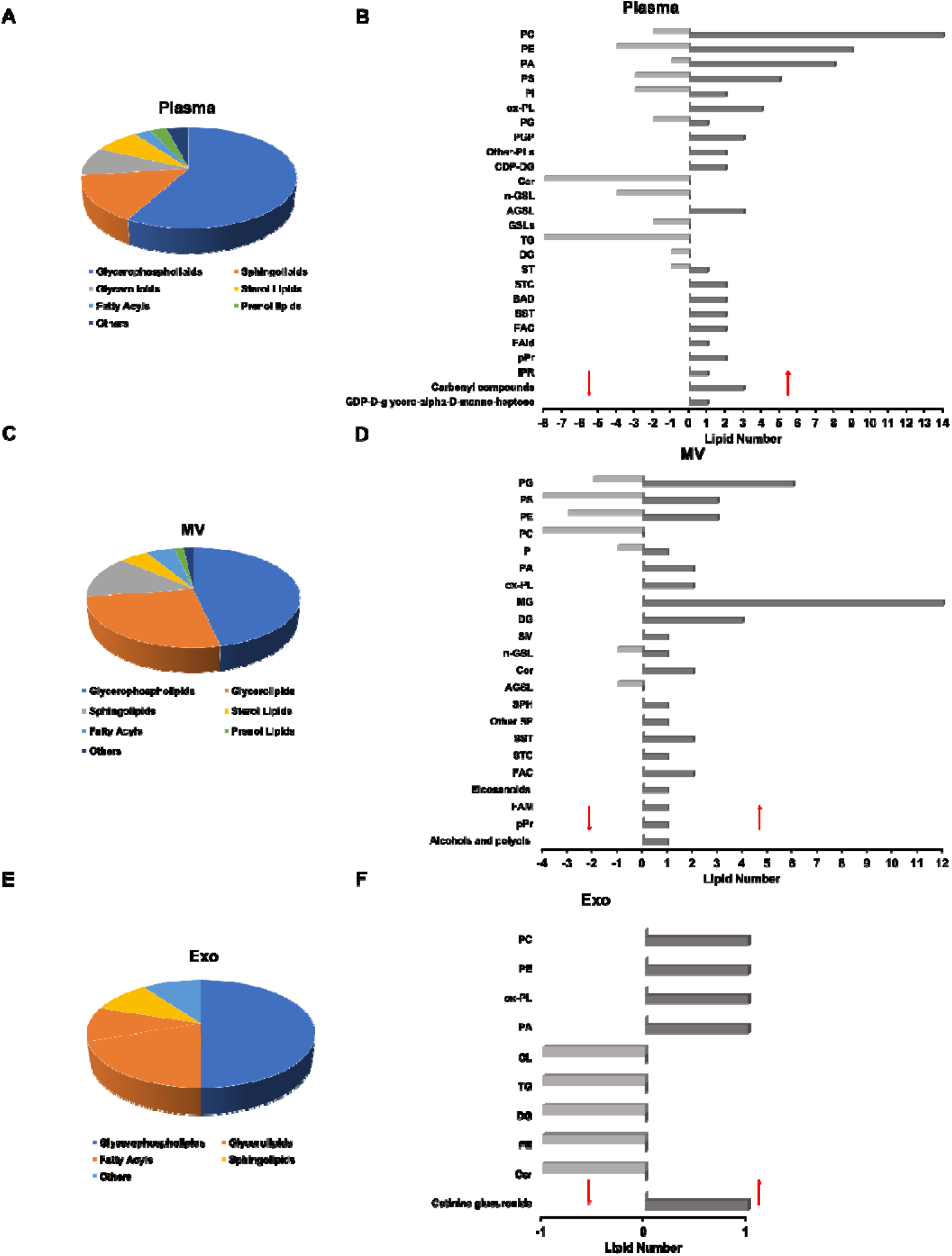
Differentially expressed lipids between preterm birth and control conditions in plasma, microvesicles and exosomes. A, C, E, a pie chart of the identified main lipid class with significant differences between the preterm birth and control conditions in plasma, microvesicles and exosomes. B, D, F, Differentially expressed lipid subclasses between the preterm birth and control conditions in plasma, microvesicles and exosomes. Upward and downward pointing arrows represent increased and decreased lipid subclasses in the preterm birth condition, respectively. MV, microvesicle; Exo, exosome.

**Table 2.** Summary of differential features and lipids between the preterm birth and control conditions.

|  | **Plasma** | **MV** | **Exo** |
| --- | --- | --- | --- |
| Differential features | 330 | 632 | 88 |
| Differential features independent<br>of other characteristics of<br>samples | 293 | 144 | 46 |
| Differential features with<br>identification | 97 | 58 | 10 |
| Candidate biomarkers | 32 | 17 | 6 |

### Application of candidate lipid biomarkers to predict the risk of preterm birth in the discovery set

To evaluate the discriminative power of lipids in plasma, MVs and Exos for the purpose of distinguishing preterm birth from the control condition, we performed ROC analysis. As shown in Table 2, 32, 17, and 6 candidate lipid biomarkers were found in plasma, MVs and Exos, respectively. Detailed information on all candidate lipid biomarkers is shown in Table S4. We further sorted candidate biomarkers by the area under the curve (AUC) values from high to low. We eventually identified five lipids with ROC > 0.8 in MVs and one in plasma (Fig. 3B). In MVs, PS (34:0), PS (O-42:0), PI (O-36:1), C24 (OH) sulfatide and PE (O-33:0) were lower in the preterm birth condition than in the full-term control condition. In plasma, OKODiA-PI was higher in the preterm birth condition than in the full-term control condition (Fig. 3A). Next, we analyzed the combined panel of these 5 lipids in MVs using the binary logistic regression rule. The AUC of the combined lipids was 0.87 with 100% sensitivity and ~ 71.2% specificity (Fig. 3C). Together, these results suggest that lipids in MVs represent the most powerful tool to distinguish preterm birth from full-term birth with sufficient sensitivity as well as specificity compared to the abilities of lipids in plasma and Exos.

**Figure 3.**
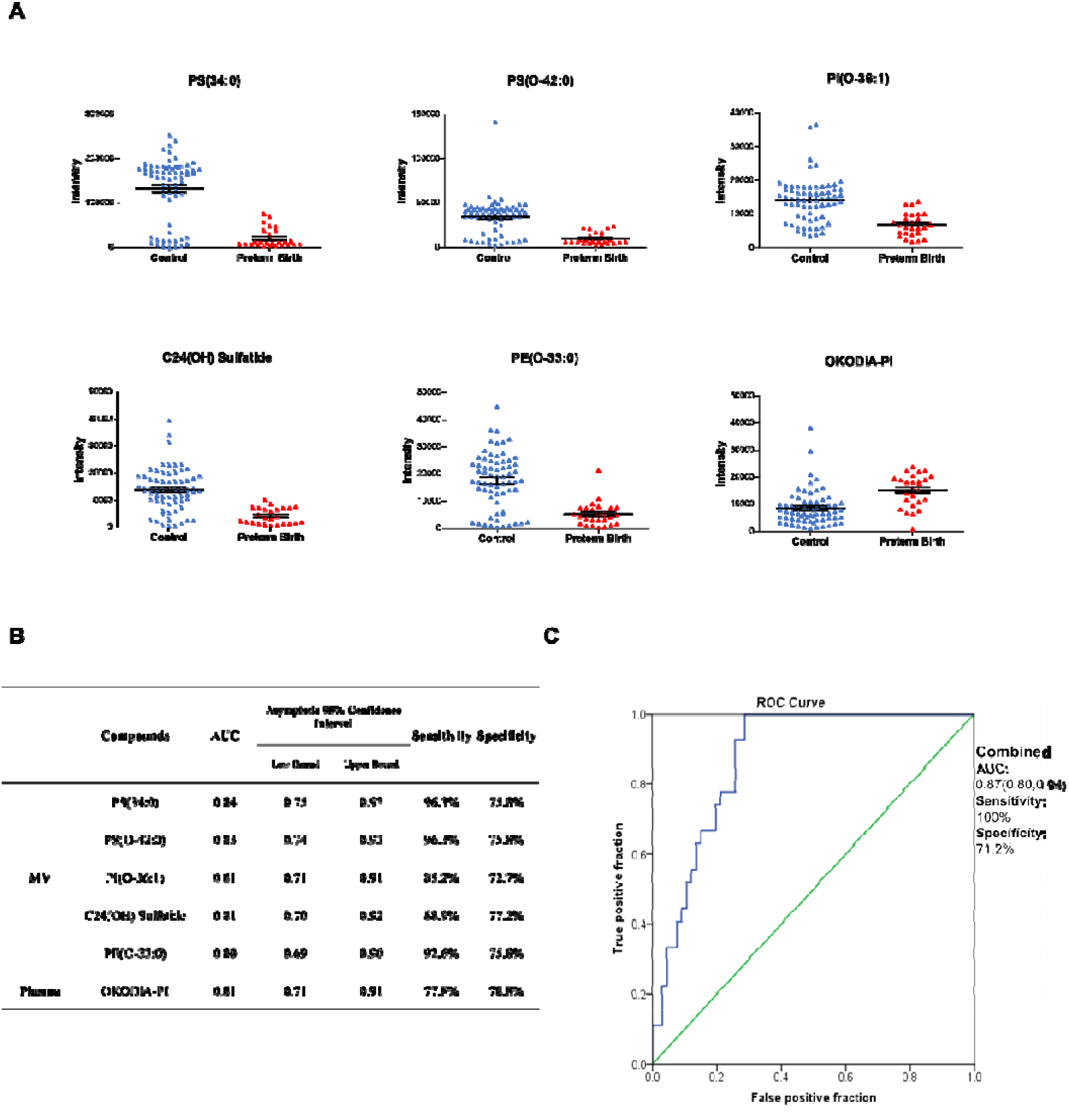
Candidate lipid biomarkers with high accuracy in the prediction of the risk of preterm birth in the discovery set. A. Scatter plot diagram of 6 candidate lipid biomarkers with significant differences between the preterm birth and control conditions. Bars represent the mean values with the standard error of the mean (SEM). B. The area under the curve (AUC), 95% confidence interval, sensitivity and specificity of the 6 lipids alone. C. The receiver operating characteristic (ROC) curve is shown for the combination of 5 lipids in microvesicles. MV, microvesicle.

### Application of candidate lipid biomarkers in MVs to predict the risk of preterm birth in the validation set

To validate these 5 candidate lipid biomarkers from MVs, an additional 83 plasma samples (41 preterm and 42 full-term deliveries) were collected (Table 1). Due to the lack of commercial standards and internal standards of these 5 lipids, we chose 4 lipids (PS (34:0), PI (O-36:1), C24 (OH) sulfatide and PE (O-33:0)) with fragments for further validation using the pseudotargeted lipidomics method [18]. The detailed MS/MS spectra are shown in Fig. S4. These 4 lipid ion pairs were constructed based on precursor ions and corresponding product ions with the highest intensity, and values of DP and CE were optimized according to corresponding lipid internal standards (Fig. 4A). Optimized values of CE and DP of lipidMix internal standards are shown in Table S5. We also applied QC samples and lipid internal standards to monitor the quality of the LC-MS system. The percentage of lipid internal standards with CV < 20% in all samples was 70% (Table S6). However, only one lipid (PS (34:0)) met the criteria (the occurrence of a lipid ion pair > 2/3 in all clinical samples and an absolute value of time deviation < 0.5 min compared with the retention time in the discovery groups). The difference in PS (34:0) was significant between preterm birth and full-term birth (P value = 0.0007). The level of PS (34:0) was lower in the preterm birth group than in the control group (Fig. 4B). We also assessed the discriminative power of PS (34:0) for distinguishing these two groups using ROC analysis. As shown in Fig. 4C, the AUC of PS (34:0) was 0.71, with 63.4% sensitivity and 76.2% specificity. The results indicate that PS (34:0) has moderate accuracy for detecting preterm birth from full-term birth in the validation set.

**Figure 4.**
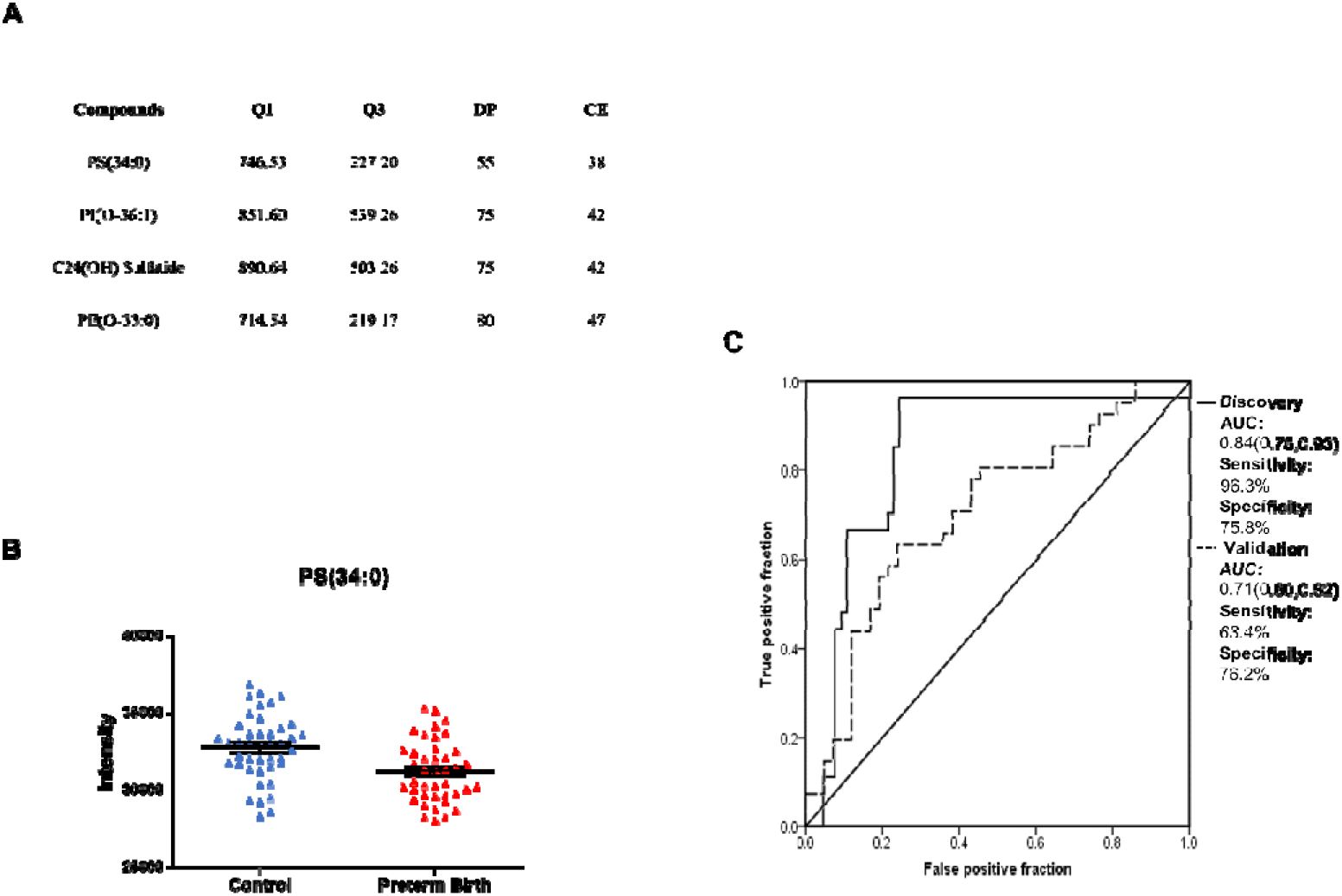
Application of candidate lipid biomarkers in microvesicles to predict the risk of preterm birth in the validation set. A. Detailed information on ion pairs of 4 candidate lipid biomarkers in microvesicles. B. Scatter plot diagram of PS (34:0) with a significant difference between the preterm birth and control conditions in microvesicles for the validation set. Bars represent the mean values with the standard error of the mean (SEM). C. The receiver operating characteristic (ROC) curve of PS (34:0) in the discovery and validation sets. The solid and dotted lines represent the discovery and validation sets, respectively.

## Discussion

In our study, we fully analyzed differential lipids associated with preterm labor in plasma, microvesicles and exosomes by performing a lipidomics profiling quantification study. We also estimated the power of 55 candidate lipid biomarkers (32, 17, and 6 candidate biomarkers in plasma, microvesicles and exosomes, respectively) to predict the risk of preterm birth. The candidate lipid biomarkers with higher accuracy in microvesicles were further validated by a pseudotargeted lipidomics approach.

Here, we used plasma, microvesicles and exosomes to investigate the changes in lipidomics associated with preterm labor. Our study showed that lipids that exhibited significant differences between preterm and full-term pregnancies in plasma, microvesicles and exosomes were different. Although we found the highest number of candidate biomarkers in plasma compared to the numbers of biomarkers in microvesicles and exosomes, the order of the power of candidate lipid biomarkers was microvesicles > plasma > exosomes. A previous study suggested that microvesicles and exosomes are potential biomarkers of metabolomics diseases [34]. The isolation of exosomes mainly based on high-speed centrifugation is time-consuming, which makes it unlikely to be suitable for clinical application [35]. However, it is more convenient to isolate microvesicles than exosomes. Our study indicates that lipids in microvesicles from plasma are ideal biomarkers for the early clinical detection of preterm birth.

In the present study, we found that DG was increased in the microvesicles of pregnant women who experienced preterm birth. DG plays an important role in the stability of the cell membrane structure and regulates the activation of protein kinase C as a second messenger [36]. The accumulation of DG contributes to systemic lipotoxicity, leading to inflammation and dysfunction [37]. Our results also show that the level of Cer is increased in microvesicles of pregnant women who experienced preterm birth. A recent study showed that circulating Cer was associated with insulin resistance and chronic low-level inflammation in individuals with obesity [38]. In our study, we show that eicosanoids are increased in the microvesicles of pregnant women who experienced preterm birth. A previous study suggested that eicosanoids played an essential role in proinflammation [38]. We also found that the level of PA is increased in microvesicles and exosomes of pregnant women who experienced preterm birth. Many studies have shown that PA is a specific activator of type I phosphatidylinositol 4-phosphate 5-kinase (PIP5K), which stimulates phosphatidylinositol (4,5) bisphosphate PI(4,5)P2 production. PA is a positive regulator of PI(4,5)P2, resulting in the accumulation of PA and PI(4,5)P2. PA and PI(4,5)P2 are necessary for vesicle exocytosis [39]. It is interesting that undecaprenol and GDP-D-glycero-alpha-D-manno-heptose are increased in the plasma of pregnant women who experienced preterm birth. We also found that undecaprenol was increased in the microvesicles of pregnant women who experienced preterm birth. Undecaprenol is involved in cell wall synthesis in gram-positive bacteria [40]. GDP-D-glycero-alpha-D-manno-heptose participates in lipopolysaccharide biosynthesis. Our study indicates that ox-PL is increased in the plasma, microvesicles and exosomes of pregnant women who experienced preterm birth. Our results suggest that LPs are also increased in the plasma, microvesicles and exosomes of pregnant women who experienced preterm birth. The accumulation of ox-PL and LPs is associated with increased oxidative stress, thus damaging the cell membrane. LPs are also important precursors and signaling molecules of inflammatory lipid mediators [41]. Our findings provide evidence that bacterial infections, chronic low-level inflammation, and oxidative stress are risk factors for preterm birth from the perspective of lipidomics. Due to a lack of automated and high-throughput tools to interpret lipidomics data, we cannot reveal the relationship between all the changes in lipids and preterm labor.

Since we could not obtain commercial standards and internal standards for the candidate lipid biomarkers of preterm birth in microvesicles, we applied a pseudotargeted lipidomics approach to validate these lipids with high accuracy. Unfortunately, only one lipid was detected in the validation set. Although lipidomics plays an important role in the occurrence and development of diseases, the current lipidomics methods are still limited. Further studies will be required to develop alternative validation methods.

In conclusion, we fully characterized the changes in lipidomics in plasma, microvesicles, and exosomes related to preterm labor. Our study suggests that bacterial infections, chronic low-level inflammation, and oxidative stress are pathophysiologic mechanisms associated with preterm labor. We also showed that altered lipids in microvesicles should allow for the early detection of preterm birth. Our insights into the differences in lipids in plasma, microvesicles and exosomes highlight the importance of lipid variants in the pathogenesis of preterm birth.

## Abbreviations

CL: Glycerophosphoglycerophosphoglycerols;
PC: Glycerophosphocholines;
PE: Glycerophosphoethanolamines;
PA: Glycerophosphates;
PS: Glycerophosphoserines;
PI: Glycerophosphoinositols;
ox-PL: Oxidized glycerophospholipids;
PG: Glycerophosphoglycerols;
PGP: Glycerophosphoglycerophosphates;
CDP-DG: CDP-glycerols;
Other-PLs: Other Glycerophospholipids;
Cer: Ceramides;
n-GSL: Neutral glycosphingolipids;
AGSL: Acidic glycosphingolipids;
GSLs: Glycosphingolipids;
SPH: Sphingoid bases;
Other SP: Other Sphingolipids;
FAC: Fatty Acids and Conjugates;
FAM: Fatty amides;
FAld: Fatty aldehydes;
FE: Fatty esters;
SM: Phosphosphingolipids;
TG: Triradylglycerols;
DG: Diradylglycerols;
MG: Monoradylglycerols;
ST: Sterols;
STC: Steroid conjugates;
BAD: Bile acids and derivatives;
SST: Secosteroids;
pPr: Polyprenols;
IPR: Isoprenoids;
LPs: Lysophospholipids;

## Acknowledgements

This work was supported by the National Key R&D Program of China under Grant number 2017YFC0908401; Key-Area Research and Development Program of Guangdong Province under Grant number 2019B020227001; and Shenzhen Municipal Government of China under Grant number JCYJ20180703093402288. The data that support the findings of this study have been deposited in the CNSA (https://db.cngb.org/cnsa/) of CNGBdb with accession code CNP0001076. We also thank the support from China National Gene Bank.

## Declaration of Interest Statemen

The authors declare that there is no conflict of interest that could be perceived as prejudicing the impartiality of the research reported.

## Supplementary material

Figure S1. TIC of QC samples.

Figure S2. PCA score plots with QC samples of plasma, microvesicles and exosomes.

Figure S3. PCA score plots, PLS-DA score plots and Volcano plots from preterm and full-term pregnant groups in plasma, microvesicles and exosomes.

Figure S4. MS/MS spectra of four candidate lipid biomarkers in microvesicles.

Table S1. The CVs of LipidMix Internal Standards in all samples in the discovery set.

Table S2. Protein concentration of plasma, microvesicles and exosomes.

Table S3. Detailed information on the differential features with identification in plasma, microvesicles and exosomes.

Table S4. Detailed information on candidate biomarkers in plasma, microvesicles and exosomes.

Table S5. The transitions of LipidMix Internal Standards.

Table S6. The CVs of LipidMix Internal Standards in all samples in the validation set.

