## Supplementary material for "Lipidomic profiling of plasma extracellular vesicles as an effective means to evaluate the risk of preterm birth": Figure S1

### Slide 1
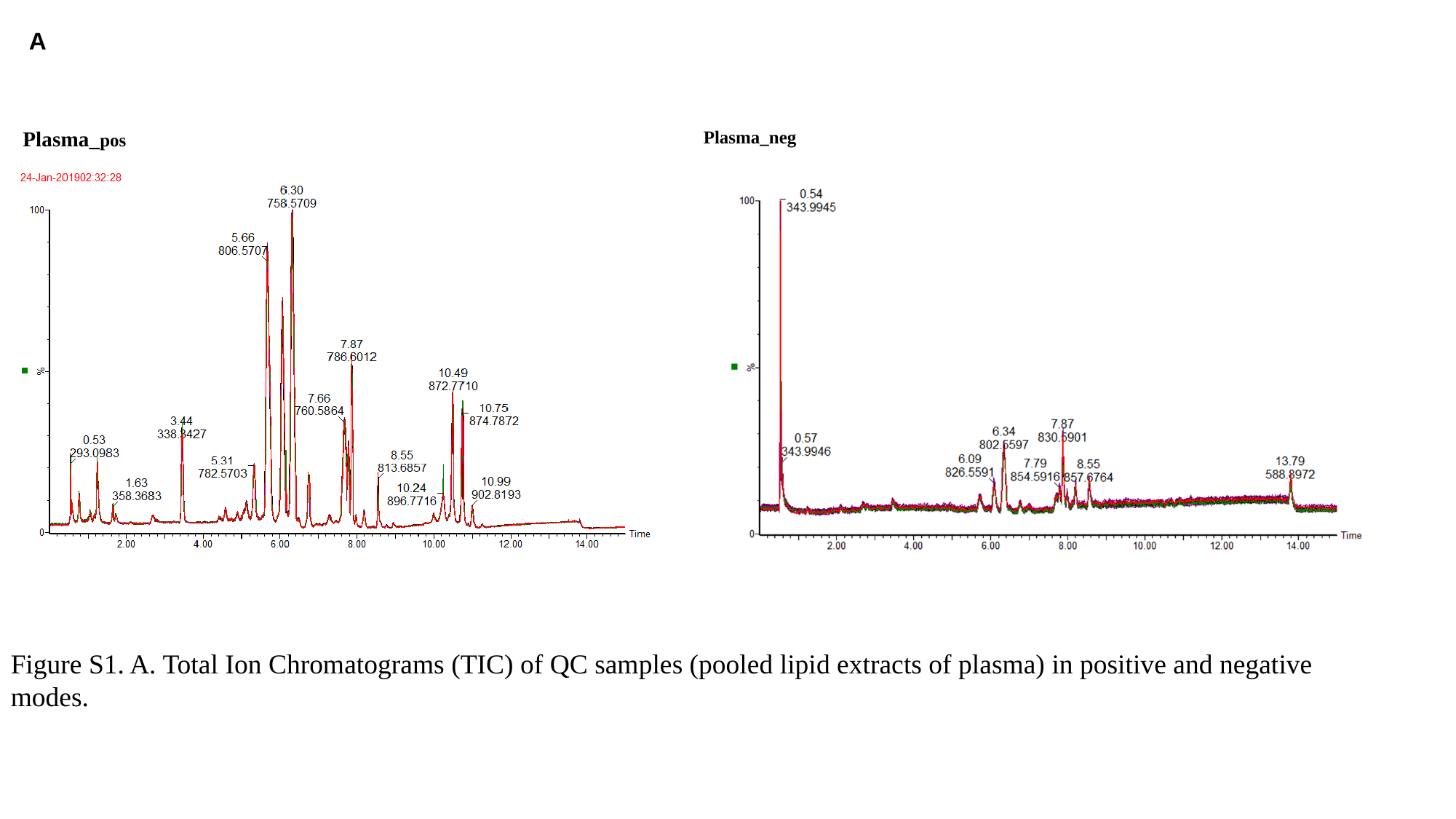

A
Plasma_pos
Plasma_neg
Figure S1. A. Total Ion Chromatograms (TIC) of QC samples (pooled lipid extracts of plasma) in positive and negative modes.

### Slide 2
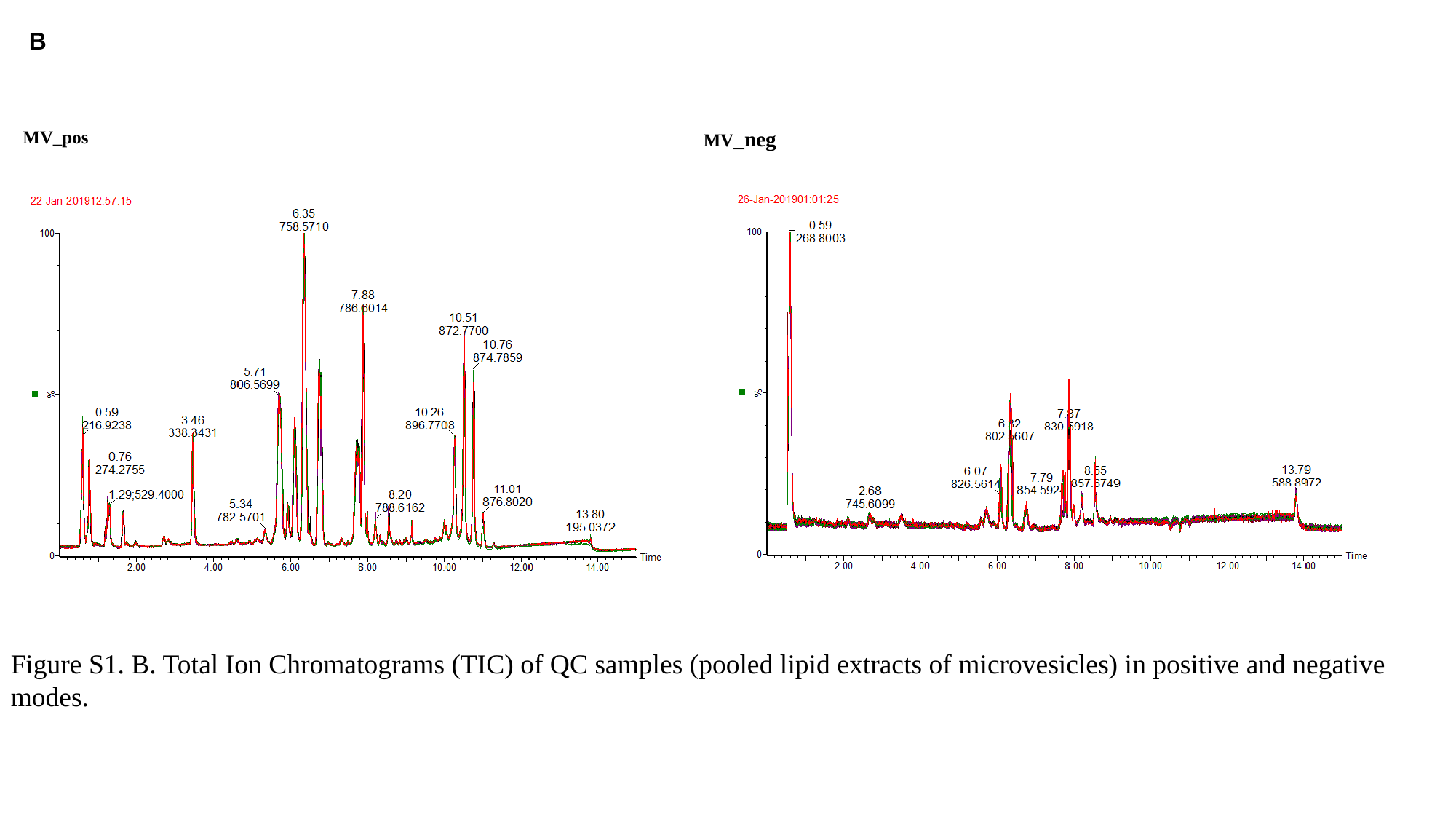

B
MV_pos
MV_neg
Figure S1. B. Total Ion Chromatograms (TIC) of QC samples (pooled lipid extracts of microvesicles) in positive and negative modes.

### Slide 3
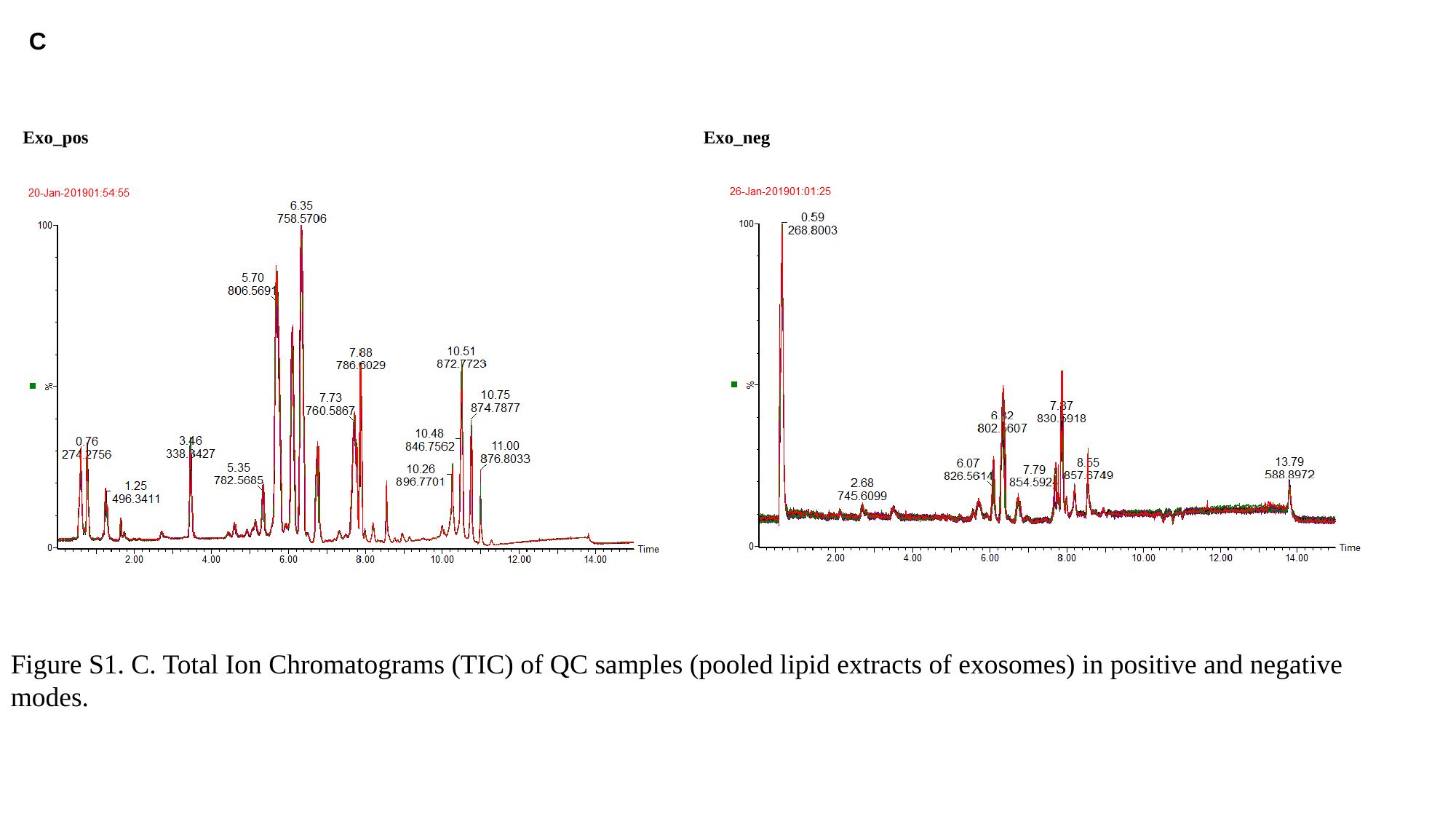

C
Exo_pos
Exo_neg
Figure S1. C. Total Ion Chromatograms (TIC) of QC samples (pooled lipid extracts of exosomes) in positive and negative modes.
