## Supplementary material for "Lipidomic profiling of plasma extracellular vesicles as an effective means to evaluate the risk of preterm birth": Figure S2

### Slide 1
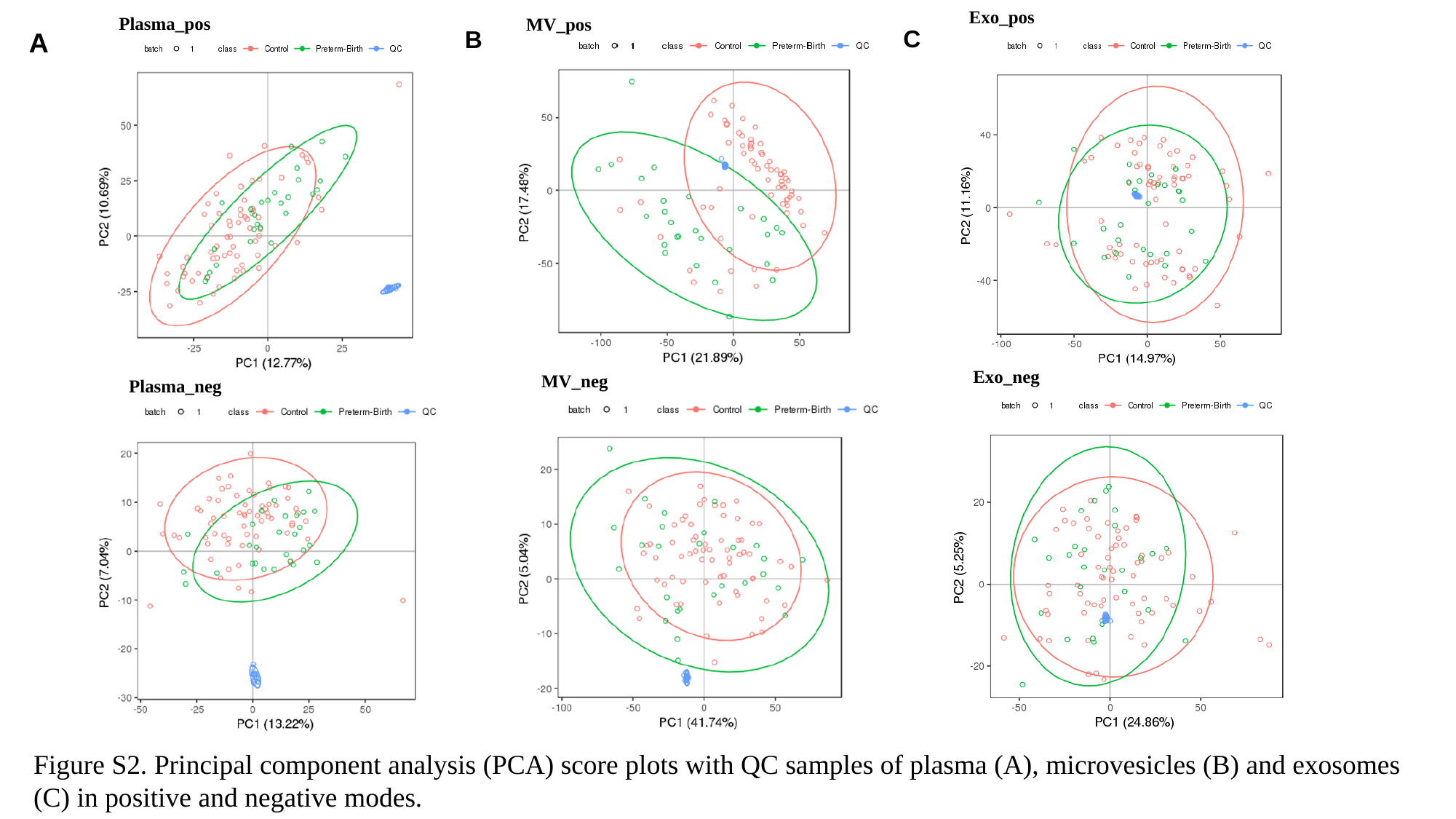

Exo_pos
Plasma_pos
MV_pos
C
B
A
Exo_neg
MV_neg
Plasma_neg
Figure S2. Principal component analysis (PCA) score plots with QC samples of plasma (A), microvesicles (B) and exosomes (C) in positive and negative modes.
