## Supplementary material for "Lipidomic profiling of plasma extracellular vesicles as an effective means to evaluate the risk of preterm birth": Figure S3

### Slide 1
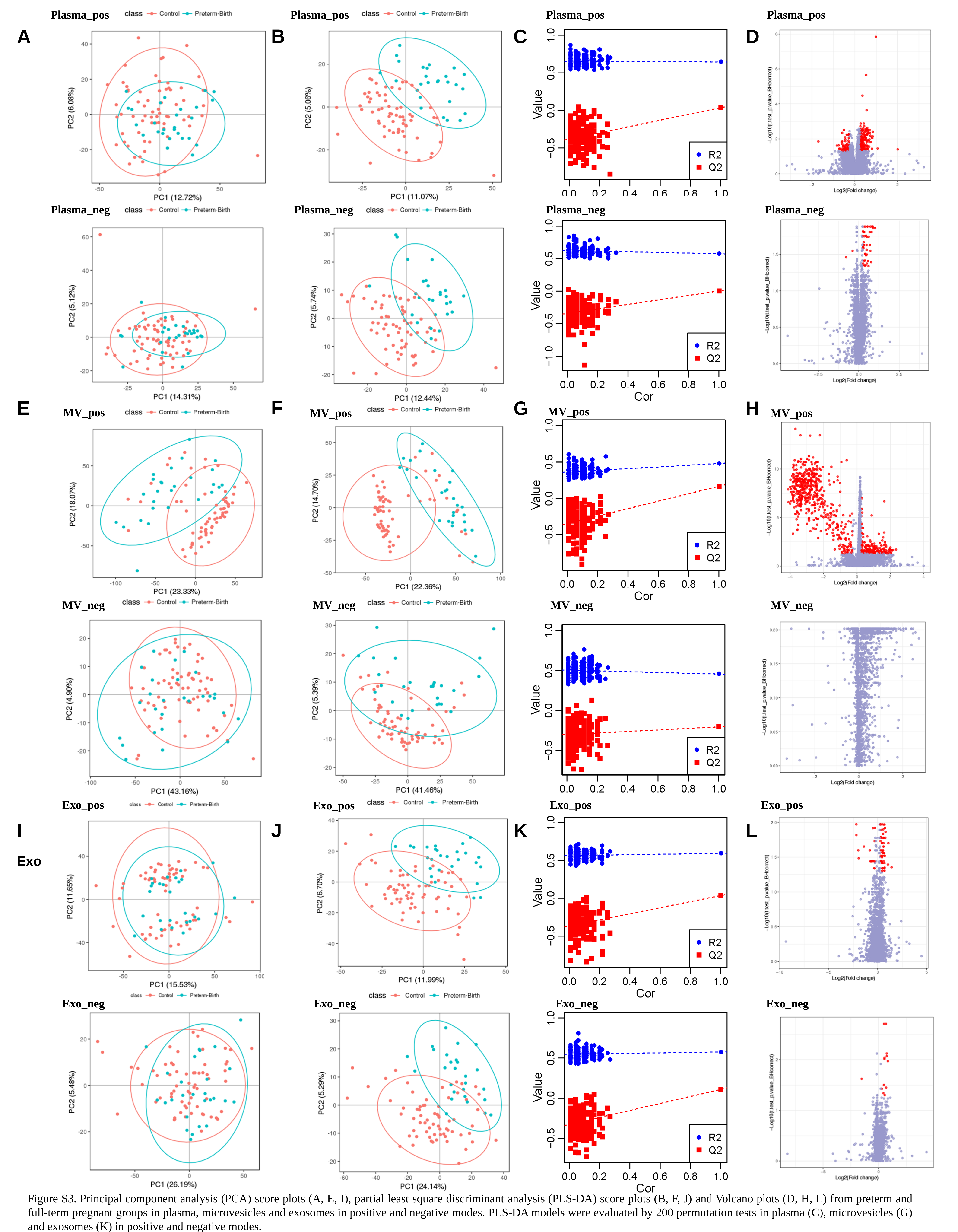

Plasma_pos
Plasma_pos
Plasma_pos
Plasma_pos
B
A
C
D
Plasma_neg
Plasma_neg
Plasma_neg
Plasma_neg
F
E
G
H
MV_pos
MV_pos
MV_pos
MV_pos
MV_neg
MV_neg
MV_neg
MV_neg
Exo_pos
Exo_pos
Exo_pos
Exo_pos
J
I
K
L
Exo
Exo_neg
Exo_neg
Exo_neg
Exo_neg
Figure S3. Principal component analysis (PCA) score plots (A, E, I), partial least square discriminant analysis (PLS-DA) score plots (B, F, J) and Volcano plots (D, H, L) from preterm and full-term pregnant groups in plasma, microvesicles and exosomes in positive and negative modes. PLS-DA models were evaluated by 200 permutation tests in plasma (C), microvesicles (G) and exosomes (K) in positive and negative modes.
