## Supplementary material for "Lipidomic profiling of plasma extracellular vesicles as an effective means to evaluate the risk of preterm birth": Figure S4

### Slide 1
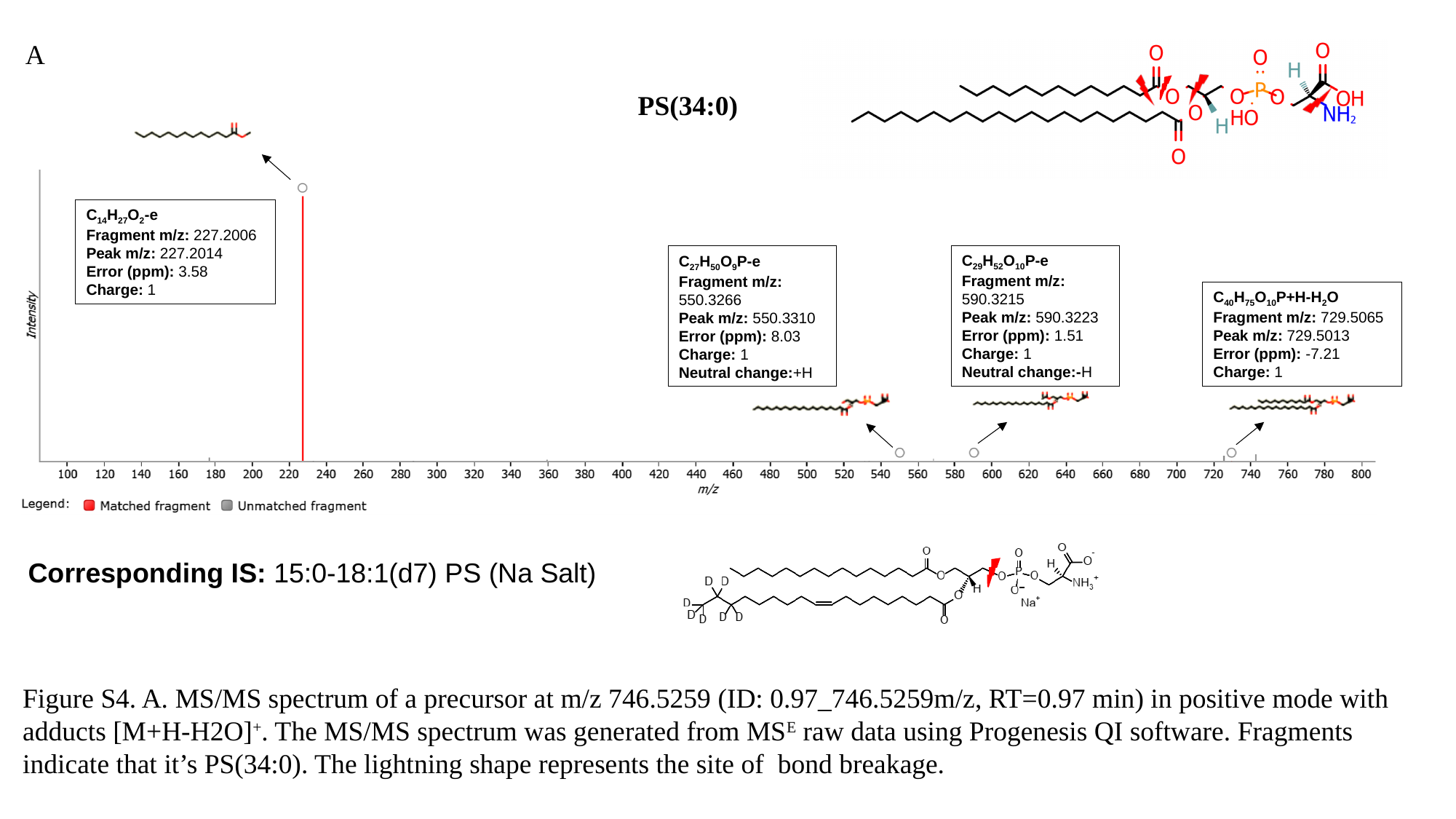

A
PS(34:0)
C_14_H_27_O_2_-e
Fragment m/z: 227.2006
Peak m/z: 227.2014
Error (ppm): 3.58
Charge: 1
C_29_H_52_O_10_P-e
Fragment m/z: 590.3215
Peak m/z: 590.3223
Error (ppm): 1.51
Charge: 1
Neutral change:-H
C_27_H_50_O_9_P-e
Fragment m/z: 550.3266
Peak m/z: 550.3310
Error (ppm): 8.03
Charge: 1
Neutral change:+H
C_40_H_75_O_10_P+H-H_2_O
Fragment m/z: 729.5065
Peak m/z: 729.5013
Error (ppm): -7.21
Charge: 1
Corresponding IS: 15:0-18:1(d7) PS (Na Salt)
Figure S4. A. MS/MS spectrum of a precursor at m/z 746.5259 (ID: 0.97_746.5259m/z, RT=0.97 min) in positive mode with adducts [M+H-H2O]^+^. The MS/MS spectrum was generated from MS^E^ raw data using Progenesis QI software. Fragments indicate that it’s PS(34:0). The lightning shape represents the site of bond breakage.

### Slide 2
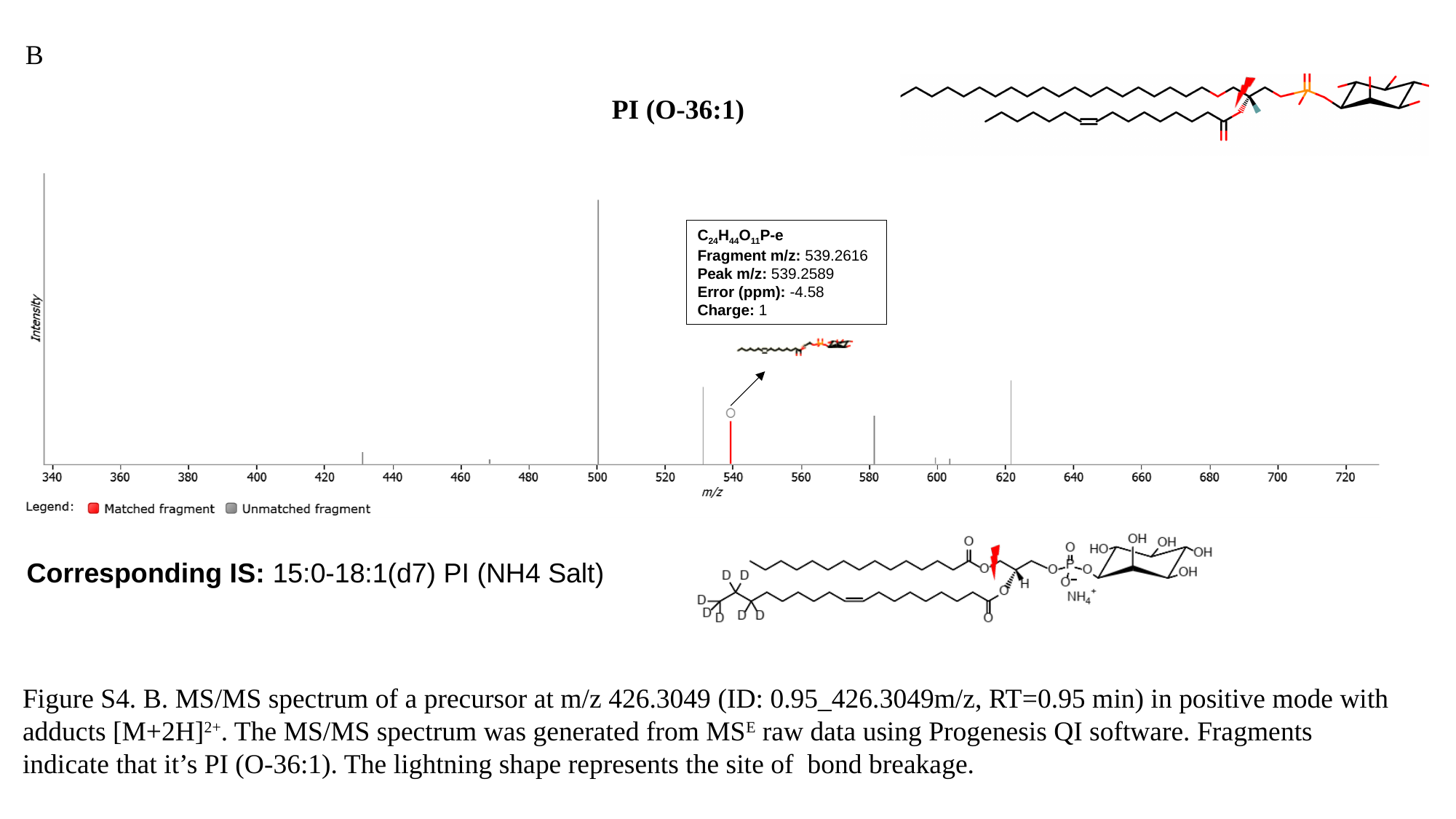

B
PI (O-36:1)
C_24_H_44_O_11_P-e
Fragment m/z: 539.2616
Peak m/z: 539.2589
Error (ppm): -4.58
Charge: 1
Corresponding IS: 15:0-18:1(d7) PI (NH4 Salt)
Figure S4. B. MS/MS spectrum of a precursor at m/z 426.3049 (ID: 0.95_426.3049m/z, RT=0.95 min) in positive mode with adducts [M+2H]^2+^. The MS/MS spectrum was generated from MS^E^ raw data using Progenesis QI software. Fragments indicate that it’s PI (O-36:1). The lightning shape represents the site of bond breakage.

### Slide 3
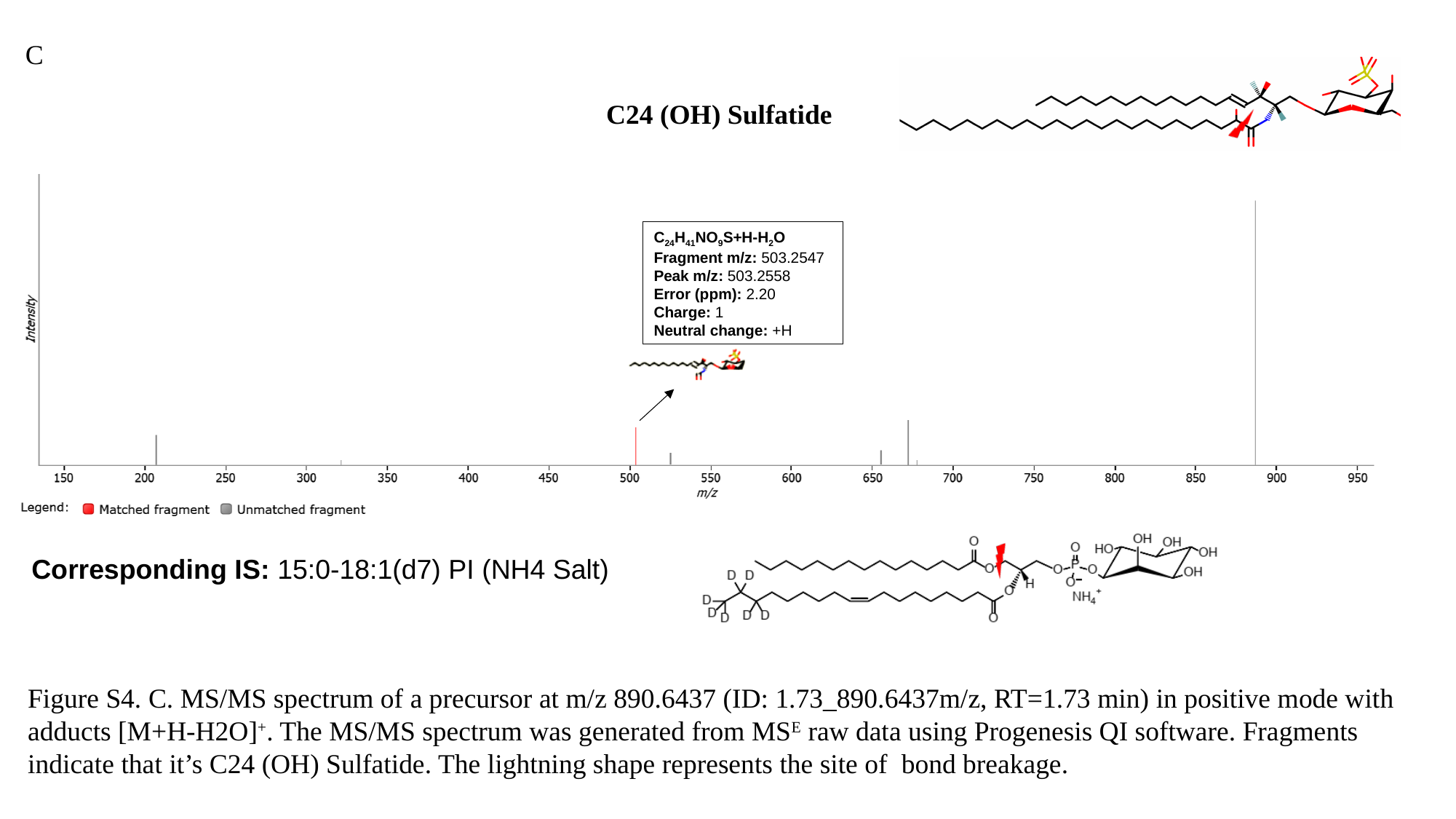

C
C24 (OH) Sulfatide
C_24_H_41_NO_9_S+H-H_2_O
Fragment m/z: 503.2547
Peak m/z: 503.2558
Error (ppm): 2.20
Charge: 1
Neutral change: +H
Corresponding IS: 15:0-18:1(d7) PI (NH4 Salt)
Figure S4. C. MS/MS spectrum of a precursor at m/z 890.6437 (ID: 1.73_890.6437m/z, RT=1.73 min) in positive mode with adducts [M+H-H2O]^+^. The MS/MS spectrum was generated from MS^E^ raw data using Progenesis QI software. Fragments indicate that it’s C24 (OH) Sulfatide. The lightning shape represents the site of bond breakage.

### Slide 4
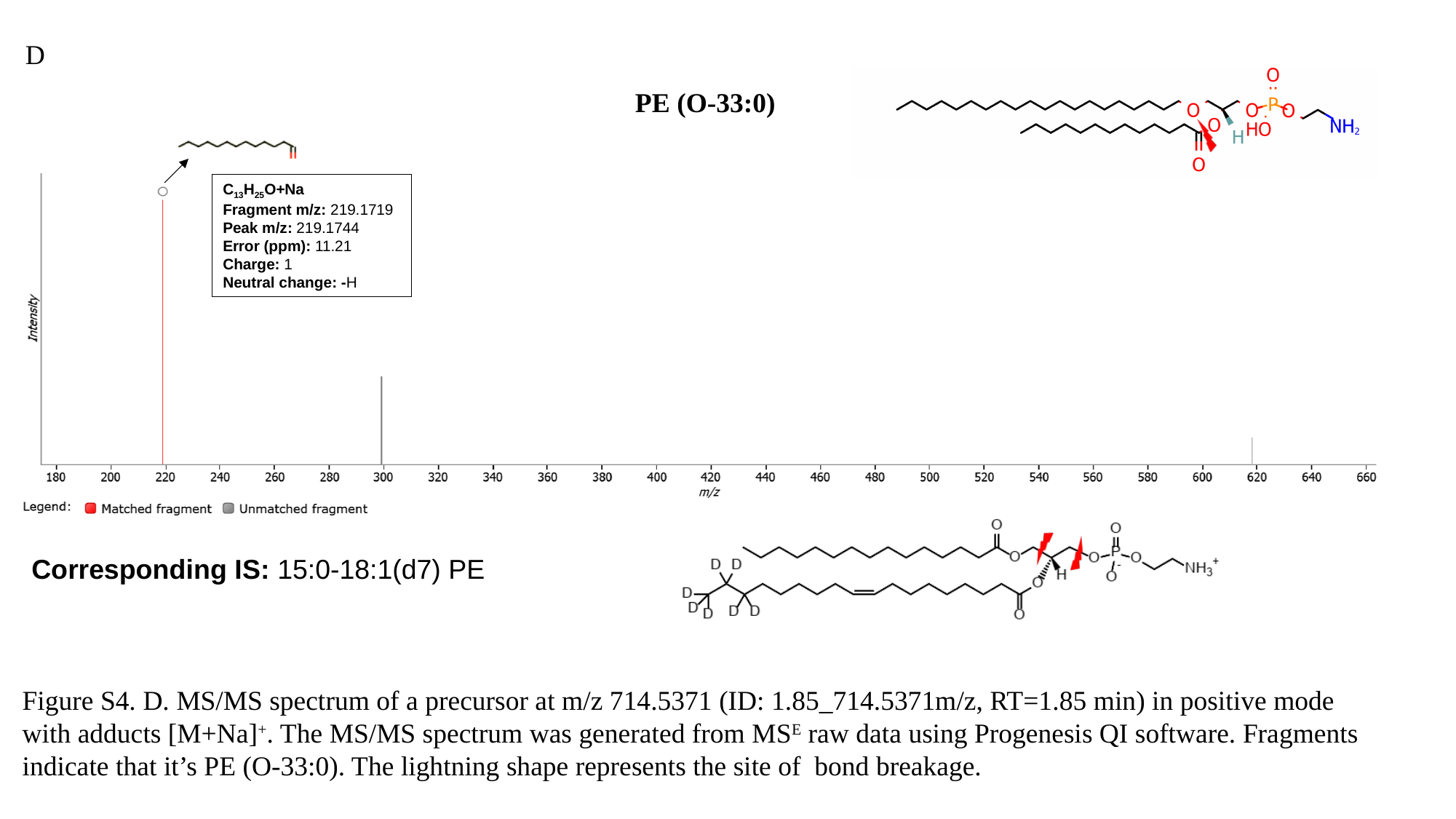

D
PE (O-33:0)
C_13_H_25_O+Na
Fragment m/z: 219.1719
Peak m/z: 219.1744
Error (ppm): 11.21
Charge: 1
Neutral change: -H
Corresponding IS: 15:0-18:1(d7) PE
Figure S4. D. MS/MS spectrum of a precursor at m/z 714.5371 (ID: 1.85_714.5371m/z, RT=1.85 min) in positive mode with adducts [M+Na]^+^. The MS/MS spectrum was generated from MS^E^ raw data using Progenesis QI software. Fragments indicate that it’s PE (O-33:0). The lightning shape represents the site of bond breakage.
